# Challenges in modelling the sediment retention ecosystem service for an ecosystem account – Examples from the Mitchell catchment

**DOI:** 10.1101/2021.08.06.455476

**Authors:** Joseph M. McMahon, Syezlin Hasan, Andrew Brooks, Graeme Curwen, Josh Dyke, Chantal Saint Ange, James C. R. Smart

## Abstract

If soil resources and the benefits derived from water quality are to be maintained, the on- and off-site effects of soil erosion must be adequately represented and communicated to decision makers so that appropriate management responses can be identified. The System of Environmental-Economic Accounting - Ecosystem Accounting (SEEA-EA), is one approach to quantify both the contributions that ecosystems make to the economy, and the impacts of economic activity on ecosystems. However, due to the difficulty of obtaining empirical data on ecosystem service flows, in many cases such quantification is informed by ecosystem service models. Previous research on the Mitchell catchment, Queensland Australia allowed us to explore the implications of using a model of hillslope erosion and sediment delivery in isolation (as represented in one of the most frequently used ecosystem service models - InVEST) by comparing such estimates against multiple lines of local empirical data, and a more comprehensive representation of locally important erosion and deposition processes through a sediment budget model. Estimates of the magnitude of hillslope erosion modelled using an approach similar to InVEST and the calibrated sediment budget differed by an order of magnitude. If an uncalibrated InVEST-type model was used to inform the relative distribution of erosion magnitude, findings suggest the incorrect erosion process would be identified as the dominant contributor to suspended sediment loads. However, the sediment budget model could only be calibrated using data on sediment sources and sinks that had been collected in the catchment through a sustained research effort. A comparable level of research investment may not be available to inform ecosystem service assessments elsewhere. The results summarised here for the Mitchell catchment demonstrate that practitioners must exercise caution when using estimates of the sediment retention ecosystem service flow which have not been calibrated and validated against locally collected empirical data.

## Introduction

Changes in land use can increase soil erosion rates, threatening agricultural sustainability and the benefits which society derives from water quality (Amundson et al., 2015; Keeler et al., 2012). If soil resources and the benefits derived from water quality are to be maintained, the on- and off-site effects of soil erosion must be adequately represented so that appropriate management responses can be identified and communicated to decision makers (Keesstra et al., 2016). The United Nations, through the System of Environmental-Economic Accounting - Ecosystem Accounting (SEEA-EA), is increasing efforts to quantify (i) the contributions that ecosystems make to the economy and society, and (ii) the impacts of economic and human activity on ecosystems (UN, 2021b). Spatially based information on ecosystem assets and the flows of services they supply to the economy and society would support decision making. However, quantifying the flow of ecosystem services, which vary spatially and temporally, to society can be challenging. Consequently, in many cases such quantification is informed by ecosystem service models (UN, 2021a).

One of the most frequently used ecosystem service models is InVEST (Integrated Valuation of Ecosystem Services and Trade-offs) (Sharp et al., 2018; UN, 2021a). Within InVEST, soil erosion and the sediment retention ecosystem service that acts to prevent soil erosion are modelled using a combination of the Revised Universal Soil Loss Equation (RUSLE), and a sediment delivery ratio (SDR) to represent the proportion of eroded sediment reaching a waterway (Sharp et al., 2018). Subsequently, all sediment that is delivered to a waterway is assumed to be transported to the catchment outlet. The RUSLE was developed to represent hillslope erosion (Wishcheimer & Smith, 1978). While InVEST documents acknowledge the importance of representing other erosion processes if they are important locally, in many locations the data or knowledge to support inclusion of these other processes in ecosystem service assessments is not available (Sharp et al., 2018).

Hillslope erosion is one of a range of different erosion processes whose magnitude can increase in response to land use change. While hillslope erosion is the dominant contributor to sediment loads in some locations (Walling & Collins, 2005), in other locations processes such as riverbank and gully erosion dominate (Olley et al., 2013)(see Figure 1 for some examples of these erosion processes). The physical configuration of a catchment influences the efficiency with which sediment contributed from various erosion processes is delivered to the catchment outlet (termed ‘sediment connectivity’)(Fryirs, 2013). Consequently, areas in which sediment can be deposited as it travels between its source and the catchment outlet can be a key consideration in evaluating this efficiency of sediment transport.

**Figure 1.**
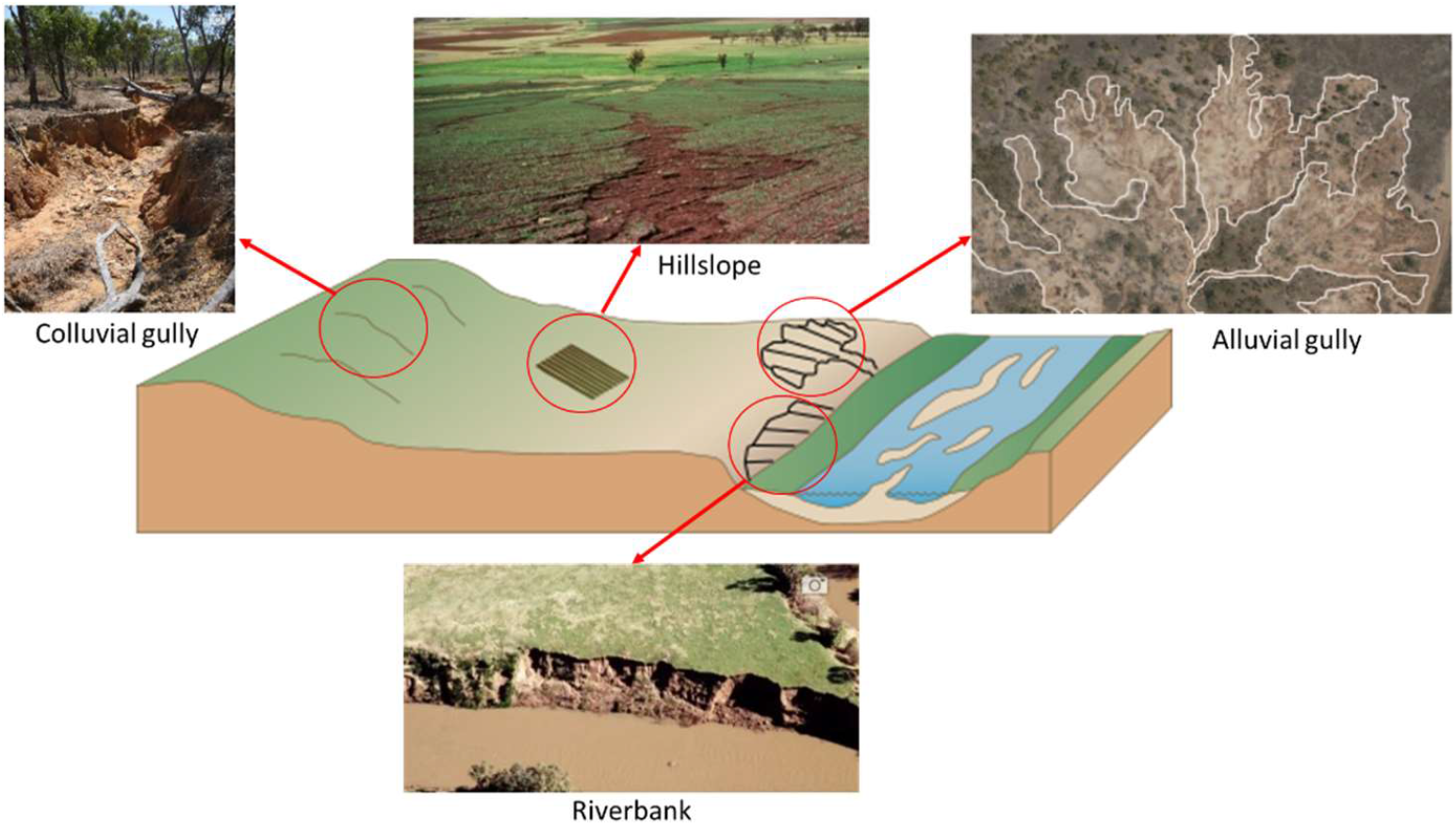
Examples of some erosion processes. Note: images of gully, hillslope and riverbank erosion are taken from Qld Government (2020), and the image of alluvial gully erosion is taken from Brooks et al. (2009)

Geomorphic systems respond dynamically to changes in erosion and deposition over time (Brierley & Fryirs, 2005; Schumm & Lichty, 1965). For example while one erosion process may be the cause of increased sediment loads historically, if this sediment has been deposited and not transported through a channel network, a different erosion process may be responsible for contemporary remobilisation of such sediment (Brierley & Fryirs, 2005; Collins & Walling, 2004; Trimble, 1983). Accurately identifying the sources and mechanisms of contemporary sediment delivery is important for determining which sections of a landscape are providing the most sediment retention and thus should be the focus for management interventions to increase supply of this ecosystem service flow.

The magnitude of different erosion processes occurring in a catchment, and the connectivity of eroded sediment to the catchment outlet, can be compiled and assessed using a sediment budget (Walling & Collins, 2008). The SedNET sediment budget model has been widely applied in Australia (McCloskey et al., 2021; Prosser et al., 2001a; Prosser et al., 2001b; Wilkinson et al., 2014; Wilkinson et al., 2009). SedNET models hillslope erosion using the RUSLE and a SDR (similar to InVEST), but also includes representations of riverbank and gully erosion, and simulates depositional areas along the channel network (Wilkinson et al., 2008). However due to the difficulty of measuring and representing various erosion processes and the efficiency of sediment transport through a channel network, sediment budgets should be viewed only as an initial hypothesis which must subsequently be tested against multiple lines of evidence (Rustomji et al., 2008).

Ecosystem services such as sediment retention are generated through the interaction of ecosystem assets at a landscape scale (UN, 2021b). Change in such ecosystem service provision over time is often evaluated with respect to a baseline condition with less intensive human modification, and contemporary conditions (note that this definition differs from that used for monetary valuation of sediment retention, which often uses bare soil as a baseline). In Australia a baseline condition of pre-European settlement is frequently used.

The aim of this article was to compare estimates of which sections of a landscape appear to be delivering the most sediment retention using: 1. The approach applied in InVEST which simulates hillslope erosion in isolation and assumes that all sediment entering a waterway is delivered to the catchment outlet; 2. Locally collected empirical data on the dominant erosion processes, locations and drivers; and 3. A sediment budget modelling approach that accounts for multiple erosion processes and depositional areas which was validated against multiple lines of locally collected empirical data. The case study area is the Mitchell catchment in northern Australia (∼72, 000 km^2^; Figure 2).

**Figure 2.**
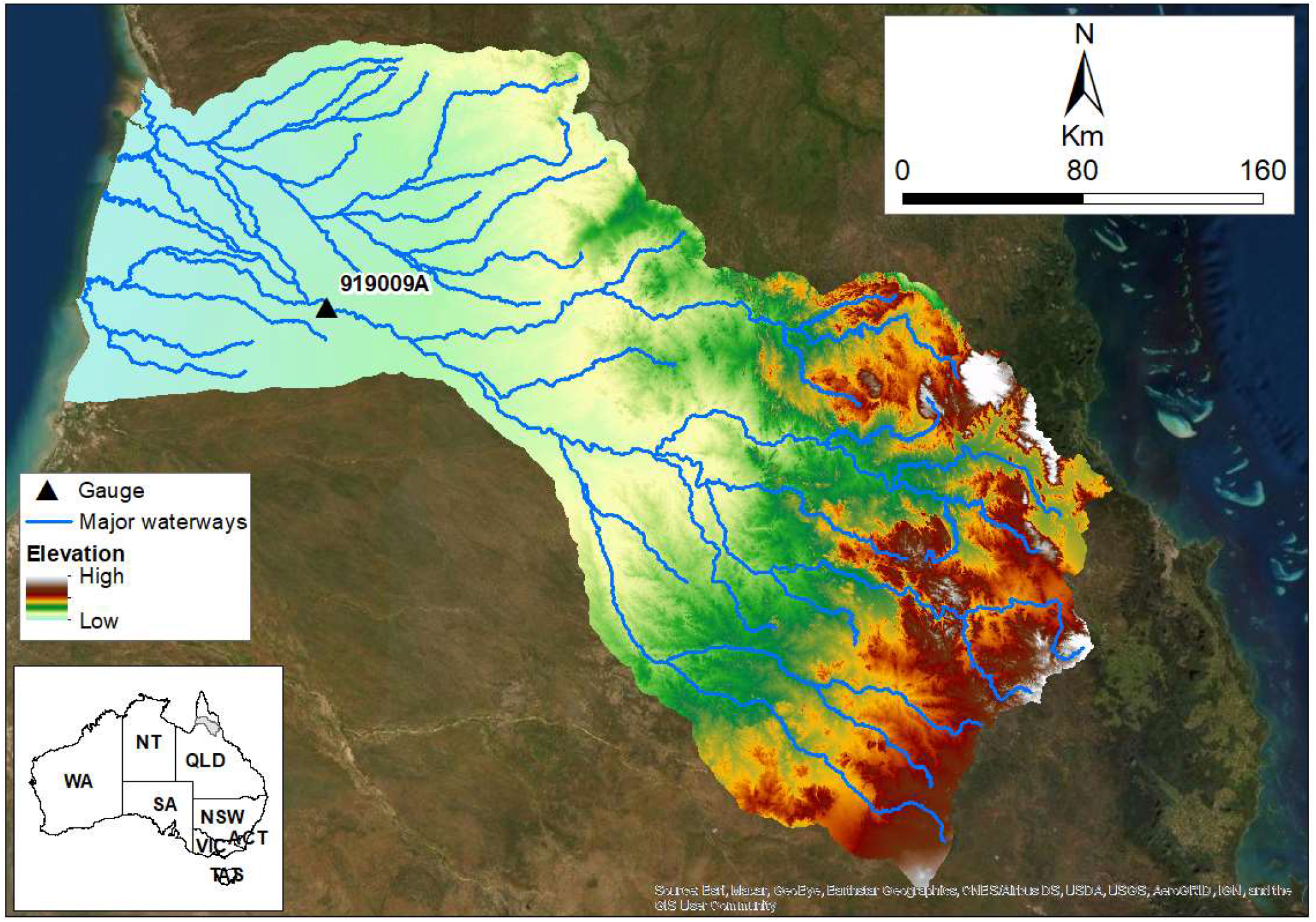
Location of the Mitchell catchment in northern Queensland, Australia.

## Regional setting

The Mitchell catchment has experienced changes to grass cover, woody vegetation, exotic species and fire regimes since European settlement (Shellberg et al., 2016). These changes have been less extensive than those in catchments in southern and eastern Australia (Brooks et al., 2009). However there are increasing calls for further development in the catchment to address growing food insecurity in Asia (Petheram et al., 2018; PMC, 2012), leading to concerns that erosion rates could increase.

The Mitchell River has one of the highest annual discharges in Australia and joins the Gulf of Carpentaria after flowing through the largest fluvial megafan in Australia (Brooks et al., 2009; Petheram et al., 2018). This megafan contains a diverse range of freshwater habitats that support the second highest fish species diversity in Australia; consequently, the catchment has a high ecological value (Petheram et al., 2018). Substantial prior research on erosion processes and sediment delivery has been undertaken in the catchment. This research has indicated:

- A particular type of gully erosion, called alluvial gully erosion (where gullies erode into alluvium adjacent to waterways as opposed to colluvial gully erosion occurring on hillsides, see Figure 1), is an important local erosion process and contributes a substantial volume of sediment to waterways (Brooks et al., 2009; Brooks et al., 2008; Shellberg et al., 2010b).
- The increase in alluvial gully erosion and its spatial distribution is influenced by both physical drivers and human related land use change (Brooks et al., 2009; Shellberg et al., 2010a; Shellberg et al., 2013b; Shellberg et al., 2016). Despite the historical human impact in the Mitchell catchment being relatively modest in comparison to catchments in southern Australia, land use changes have increased the magnitude and distribution of alluvial gully erosion substantially (Shellberg et al., 2016).

Prior research created a sediment budget for the catchment using the SedNET framework (Rustomji et al., 2010). The Rustomji et al. (2010) report describing the creation of the sediment budget details the erosion estimates from an initial model and subsequent iterations in which the model was progressively calibrated against multiple lines of evidence and incorporated important local erosion processes such as alluvial gully erosion. The sediment budget calibrated against contemporary data was subsequently used to estimate sediment loads prior to European settlement. The detail provided in Rustomji et al’s report allows a comparison to be made between the InVEST modelling approach, which estimates sediment contributed from hillslope erosion in isolation and assumes all of that sediment is transported to the catchment outlet, and a sediment budget modelling approach whose predictions have been validated against multiple lines of evidence.

## Method

### Hillslope erosion

Hillslope erosion was modelled within the Rustomji et al. (2010) report using a national grid of mean hillslope erosion rate (Lu et al., 2003), calculated using the RUSLE (Renard et al., 1997). An initial estimate of the sediment delivery ratio (SDR) was calculated using mean sub-catchment slope and mean sub-catchment foliage projective cover (Armston et al., 2004) as a measure of vegetation-related hydraulic roughness, and it was subsequently adjusted in relation to the empirical data.

### Locally collected empirical data on the dominant erosion processes and locations

Several empirical datasets were available in the Mitchell catchment to evaluate the initial hillslope erosion estimates. These data described: (i) the relative distribution of erosion processes, (ii) the magnitudes of erosion processes in sub-catchments, and (iii) suspended sediment load in absolute terms (i.e. tonnes year^-1^).

Evaluation of the relative distribution of erosion processes was informed by radionuclide analyses (Caitcheon et al., 2011; Caitcheon et al., 2012). By comparing naturally occurring and anthropogenic fallout radionuclides in sediment samples collected in the catchment, it is possible to determine whether the sample sediment came from surface or subsurface erosion (Olley et al., 1993; Wallbrink & Murray, 1993). In relation to the erosion processes outlined above and in Figure 1, only hillslope erosion would have a surface erosion fallout radionuclide signature.

Evaluation of the magnitude of sub-catchment erosion was informed by geochemical fingerprinting analyses (Caitcheon et al., 2011). Such analyses can link sampled sediment to parent geologies and therefore provide an independent check on predicted erosion rates in sub-catchments with unique geochemical signatures.

Finally, suspended sediment samples were collected by the Queensland Government at several gauge locations throughout the catchment and were used together with measured discharge data to estimate suspended sediment load (Rustomji et al., 2010). Despite some limitations in terms of data collection (e.g. width and depth integrated samples were not available), suspended sediment load data were compared to other estimates from Australia (Wasson, 1994) and found to be sufficiently robust (Rustomji et al., 2010).

### Sediment budget modelling approach

The sediment budget methodology employed by Rustomji et al. (2010) used the SedNET algorithms as a starting point and incorporated locally specific data and parameters where appropriate and/or available. The influence of hillslope erosion, both colluvial and alluvial gully erosion, riverbank erosion, and floodplain deposition on suspended sediment (sediment <63 µm) load was modelled at a sub-catchment resolution (Figure 3). The model was validated using the radionuclide, geochemical and suspended sediment load empirical data outlined above (Rustomji et al., 2010). These data have proven valuable for calibration of sediment budgets elsewhere (Rustomji et al., 2008; Walling & Collins, 2008). Bed material (sediment >63 µm) transport was also modelled in Rustomji et al. (2010); however, as bed material transport did not link explicitly to suspended sediment aspects of the model, and bed material transport is not simulated in InVEST, it is not discussed further here. The data and approach applied by Rustomji et al. (2010) for modelling erosion processes, in addition to hillslope erosion, and deposition are summarised in the following section.

**Figure 3.**
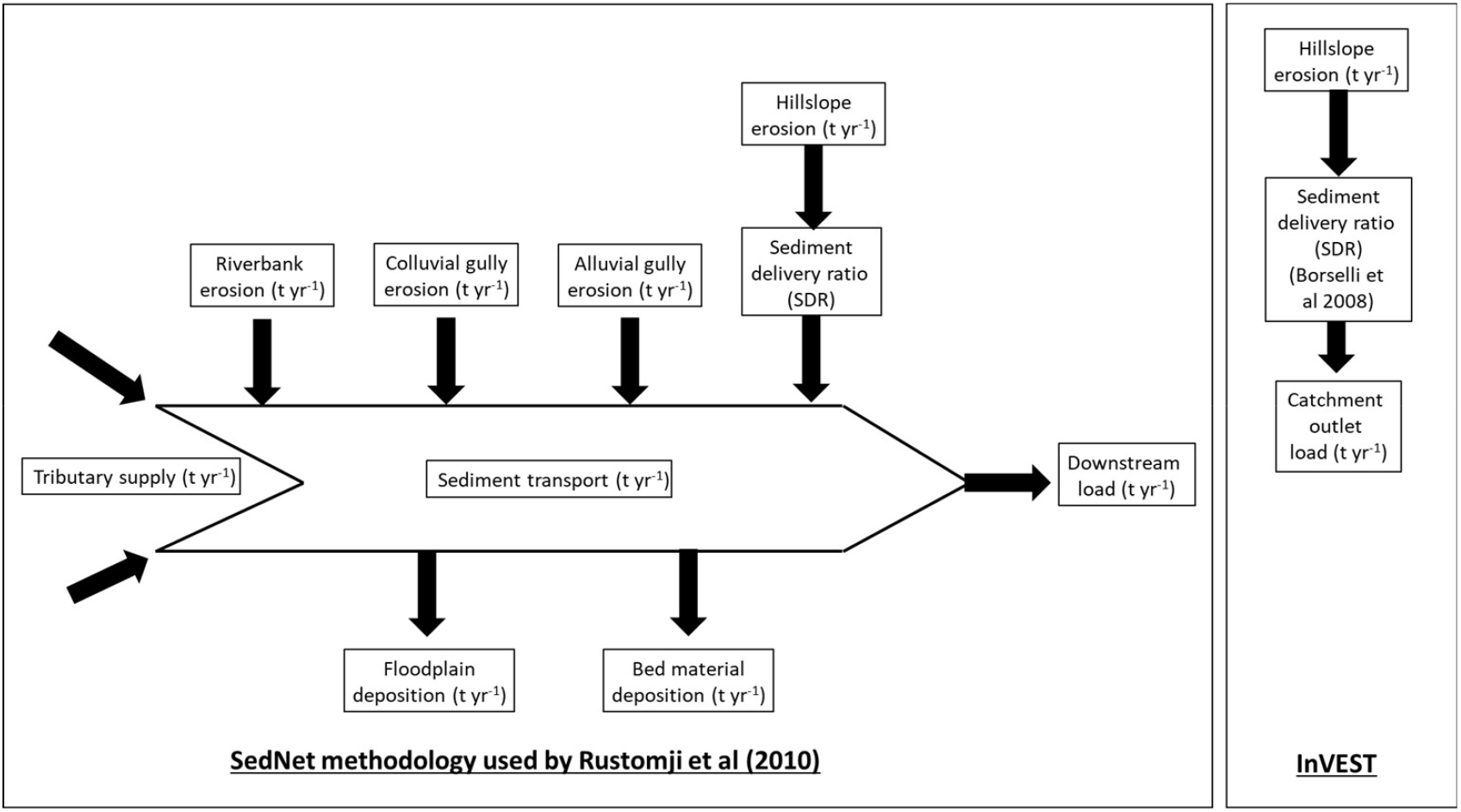
Components of the sediment budget modelled by Rustomji et al. (2010) using a modified SedNET approach, and InVEST (Sharp et al., 2018)

#### Colluvial gully erosion

Colluvial gully erosion was modelled using national data on colluvial gully density (Hughes et al., 2001) and the standard approach to modelling gully erosion within SedNET (Prosser et al., 2001a). Gullies were assumed to have initiated 100 years ago in response to land use change associated with European settlement, and were currently eroding at a lower rate (Rustomji et al., 2010).

#### Alluvial gully erosion

The extent of alluvial gully erosion was mapped using remotely sensed data by Brooks et al. (2008) and Brooks et al. (2009). Alluvial gullying extent was then combined with estimates of median gully retreat rates calculated from aerial photos and field data by Shellberg et al. (2010b), to derive an estimate of alluvial gully erosion in tonnes year^-1^. These estimates were then expressed in kilometres of gully per kilometre squared of land area to be consistent with other SedNET equations (Rustomji et al., 2010). Based on data presented in Shellberg et al. (2010b), and in contrast to colluvial gullies, alluvial gullies appeared to be actively eroding and were represented as such in the model (Rustomji et al., 2010).

#### Riverbank erosion

Riverbank erosion within the stream network in each sub-catchment was modelled using the methodology presented in Wilkinson et al. (2006) (Rustomji et al., 2010). This methodology required streampower (the product of bankfull discharge and slope), the proportion of woody riparian vegetation cover estimated using foliage projective cover, the proportion of flat alluvium derived using the methods of Gallant and Dowling (2003), sub-catchment waterway length, and riverbank height derived from field surveyed cross sections as well as cross sections extracted from a 30m Digital Elevation Model (DEM). Bankfull discharge was estimated at several locations throughout the catchment, and a predictive model was created to estimate it for all required sub-catchments.

#### Floodplain deposition

Data on floodplain inundation area were derived from an approximate 278m DEM (Pickup & Marks, 2001). For a series of cross sections, peak flood discharge (m^3^ s^-1^) was modelled using upstream contributing area and mean upstream catchment rainfall (Rustomji, 2010). Peak flood discharge was compared to bankfull discharge for each sub-catchment to estimate the median overbank flow. Floodplain deposition was calculated using median overbank flow, predicted sediment load, and an estimate of how long sediment-laden water remains on the floodplain (Rustomji et al., 2010).

### Pre-European settlement erosion rates

The erosion process models (e.g. hillslope, riverbank, gully etc) of Rustomji et al’s final calibrated sediment budget model were adjusted, where appropriate, to represent pre-European settlement conditions, erosion rates and suspended sediment yield. As minimal woody riparian vegetation clearing was assumed to have occurred in the catchment after European settlement, the riverbank erosion model was not modified for pre-European settlement conditions (Rustomji et al., 2010). However, hillslope erosion estimates were adjusted to account for the introduction of cattle grazing, agriculture and vegetation clearing. For colluvial gully erosion, the assumption regarding the change in erosion rate over time in response to European settlement land use change had not been verified and therefore it was modelled as for current conditions and assumed to be broadly representative of pre-European conditions. Based on research linking alluvial gully erosion to land use changes associated with European settlement (Shellberg et al., 2010a; Shellberg et al., 2010b), alluvial gully erosion for the pre-European sediment budget was set to 5% of current rates (Rustomji et al., 2010). Two separate pre-European condition sediment budgets were created; the first budget only included changes related to hillslope erosion, whereas the second budget included changes to both hillslope and gully erosion (note that floodplain deposition was included in both budgets).

## Results

The sediment budget model created by Rustomji et al. (2010) initially indicated that hillslope erosion contributed around 60% of the predicted suspended sediment yield (Table 1). Hillslope erosion hotspots were identified in the upper sections of the catchment where slopes were relatively high, and foliage projective cover was relatively low. The predicted magnitude of hillslope erosion was considered to be unrealistically high, particularly in steep sections of the landscape with minimal vegetation cover. For example, the RUSLE implicitly assumes that hillslope erosion rate increases with slope. However, in many steep locations there may not be a large volume of soil available for erosion (Lu et al., 2003). If these steep areas also have minimal vegetation cover, this can result in unrealistically high predicted hillslope erosion rates from the RUSLE. In the first iteration of Rustomji et al’s sediment budget model the RUSLE was predicting high hillslope erosion rates due to relatively high slope and relatively low vegetation cover. Therefore, in the second model iteration hillslope erosion rate was capped at 50 tonnes hectare^-1^ year^-1^.

**Table 1.** Erosion and deposition process contributions to suspended sediment load at the Mitchell catchment outlet for different calibration iterations from Rustomji et al. (2010). All values are mean annual rates (i.e. rates per year). Note that gully input includes both colluvial and alluvial gully sources and that data for the second iteration were not presented in Rustomji et al’s report.

| *Model iteration* | *Hillslope input (kt yr<sup>-1</sup>)* | *Gully input (kt yr<sup>-1</sup>)* | *Riverbank input (kt yr<sup>-1</sup>)* | *Floodplain deposition (kt yr<sup>-1</sup>)* | *Suspended sediment load (kt yr<sup>-1</sup>)* |
| --- | --- | --- | --- | --- | --- |
| First | 8138 | 4144 | 4987 | 3542 | 13726 |
| Second | - | - | - | - | - |
| Third | 1067 | 4144 | 4987 | 1476 | 8722 |
| Fourth | 1067 | 4144 | 2493 | 1247 | 6458 |
| Fifth | 892 | 4144 | 1247 | 3364 | 2919 |

Empirical radionuclide data suggested that hillslope erosion was a minor sediment source in the catchment, and measured suspended sediment load data in the catchment indicated that the second model iteration was over predicting suspended sediment load. Consequently, the hillslope SDR was reduced from a mean value of 5% to a mean value of 1%. Subsequently, the predicted hillslope erosion became a minor sediment source in the catchment.

The predicted suspended sediment load from the third model iteration was still closer to the maximum observed suspended sediment load in the catchment – rather than the mean load (see Table 1 and Table 2). Out of the remaining erosion processes, Rustomji et al had greater confidence in the estimates for alluvial gully erosion as this process was characterised by local empirical data. Therefore, a calibration parameter in the riverbank erosion model was adjusted to reduce the contribution from riverbank erosion in the next sediment budget model iteration.

**Table 2.** Summary statistics for annual (based on water year) suspended sediment load (kt year^-1^) estimates derived from observed data as presented in Rustomji et al. (2010)

| *Gauge* | *Minimum* | *1<sup>st</sup> quartile* | *Median* | *Mean* | *3<sup>rd</sup> quartile* | *Maximum* |
| --- | --- | --- | --- | --- | --- | --- |
| 919009A | 228 | 766 | 1455 | 2079 | 2603 | 8986 |

Reducing the contribution from riverbank erosion changed the modelled relative proportions of surface and subsurface sediment to a degree which was not supported by the radionuclide data. Additionally, the predicted suspended sediment load was still too high when compared to the observed suspended sediment load (see Table 1 and Table 2), particularly in the lower catchment (note sub-catchment loads informed by geochemical analyses are not presented here). Consequently, in the final model iteration hillslope erosion was capped at 35 tonnes hectare^-1^ year^-1^, the riverbank erosion coefficient was reduced further, and floodplain deposition was increased.

The final model predicted a total suspended sediment yield of 2919 kt year^-1^ at the catchment outlet (Table 1). At the 919001A gauge (Figure 1), model predictions (2814 kt year^-1^) agreed reasonably well with the third quartile of the observed suspended loads (Table 2). The majority of the suspended sediment load predicted by the sediment budget model was contributed by alluvial gully erosion (Table 1 – fifth iteration)(the breakdown of contributions from colluvial and alluvial gully erosion is not presented here). Hillslope erosion was a minor contributor, and 3364 kt yr^-1^ of sediment was deposited on floodplains.

### Comparing the results to the InVEST modelling approach

As a reminder, the InVEST modelling approach estimates sediment contributed from hillslope erosion in isolation and assumes all of that sediment is transported to the catchment outlet. In contrast, the sediment budget modelling approach simulates multiple erosion processes and sediment deposition on floodplains (see Figure 3). The approach to modelling *hillslope erosion* used by Rustomji et al. (2010) differs from that suggested in InVEST only in its calculation of the SDR. While different methods are used for this calculation (for example the InVEST SDR approach is calculated using the formula suggested in Borselli et al. (2008)), differences in the SDR methodology are unlikely to explain the need for the model iterations to align model predictions with multiple lines of locally collected empirical evidence. Acknowledging these methodological differences, the model results presented above can be used to compare the implications of using a hillslope erosion and SDR estimate in isolation of locally collected empirical data, in contrast to a more comprehensive sediment budget.

The first iteration of the Rustomji et al. (2010) model would have mistakenly identified that hillslope erosion was the main erosion process in the catchment and mis-identified the upper catchment was the highest eroding location in the catchment. As floodplain width, and consequently floodplain deposition, generally increased in a downstream direction a proportion of sediment contributed from hillslope erosion is deposited on floodplains downstream. If floodplain deposition was not represented, as is the case in InVEST, the contribution from hillslope erosion to suspended sediment yield at the catchment outlet would thus be even higher. The data on alluvial gully erosion collected in previous studies was crucial for characterising and representing this process in the sediment budget model. These alluvial gullies were actively eroding and delivered the bulk of eroded sediment directly to waterways due to their location directly adjacent to waterways. Similarly, the radionuclide and geochemical data were crucial for confirming, respectively, that sub-surface sources were the dominant contributor to suspended sediment loads in the Mitchell, and that sub-catchment loads predicted by the model were supported by observations.

### Impact of modelling approaches on estimated erosion rate before European settlement

The initial pre-European settlement sediment budget which evaluated changes in hillslope erosion only, estimated that hillslope erosion contributed approximately 50% less sediment before European settlement (Table 3). However, floodplain deposition would still have been occurring so the pre-European suspended sediment yield at the catchment outlet was only predicted to be 3% lower than the load predicted for current conditions by the final iteration of Rustomji et al.’s sediment budget model (Table 1 – fifth iteration). Conversely when changes to gully erosion rate were included in the second iteration of the pre-European settlement sediment budget, Rustomji et al. (2010) suggest that suspended sediment load at the catchment outlet had approximately doubled since European settlement.

**Table 3.** Estimates from two iterations of the sediment budget model representing pre-European settlement conditions presented in Rustomji et al. (2010).

| *Model iteration* | *Hillslope input*<br>*(kt yr<sup>-1</sup>)* | *Gully input*<br>*(kt yr<sup>-1</sup>)* | *Riverbank input*<br>*(kt yr<sup>-1</sup>)* | *Floodplain deposition*<br>*(kt yr<sup>-1</sup>)* | *Suspended sediment load*<br>*(kt yr<sup>-1</sup>)* |
| --- | --- | --- | --- | --- | --- |
| First iteration | 406 |  |  | 2956 | 2840 |
| Second iteration | 406 | 647 |  | 1248 | 1052 |

## Discussion

### Implications for sediment retention modelling in the Mitchell catchment

Estimates of the magnitude of hillslope erosion modelled using an approach similar to that in InVEST (which simulates hillslope erosion in isolation and does not account for depositional areas downstream) and a more detailed sediment budget informed and calibrated by local empirical data, differed by an order of magnitude in the Mitchell catchment (for example 8138 kt year^-1^ for the first iteration and 892 kt year^-1^ in the final iteration, Table 1). Studies comparing modelled and empirical hillslope erosion rates in a catchment adjacent to the Mitchell catchment, suggest that the modelled hillslope erosion rates in the calibrated sediment budget model may still be overestimated (Brooks et al., 2014). If an uncalibrated InVEST model was used to inform the *relative* distribution of erosion magnitude and significance, as suggested by Sharp et al. (2018), the results presented indicate the approach would not correctly identify the dominant erosion process contributing to suspended sediment loads in the Mitchell catchment. However, this could only be ascertained, and the final calibrated sediment budget model developed, using the considerable quantity of empirical data on sediment sources and sinks that had been collected in the catchment through a sustained and concerted effort from multiple researchers over many years (Brooks et al., 2009; Brooks et al., 2008; Caitcheon et al., 2011; Caitcheon et al., 2012; Shellberg et al., 2010a; Shellberg et al., 2013a; Shellberg et al., 2010b; Shellberg et al., 2016). A comparable level of research investment may not be available to inform ecosystem service assessments in catchments elsewhere. If this is the case, ecosystem service predictions from model results should be used with extreme caution.

A major finding of both the local empirical data and the final sediment budget model calibrated against those data was that alluvial gully erosion was the dominant contributor to end of catchment suspended sediment load in the catchment. The majority of the data that informed this estimate were obtained from the mid to lower sections of the catchment adjacent to waterways. Alluvial gully erosion is also known to occur in agricultural areas in the upper catchment, however no data were available to assist quantification of alluvial gully sources from these locations in the sediment budget (Rustomji et al., 2010). It appears as though soil characteristics and climatic conditions predispose the catchment to alluvial gully erosion (Brooks et al., 2009; Shellberg et al., 2016). While all sections of the catchment appear prone to this type of erosion, erosion appears more severe in the deeper alluvial soils in the mid to lower catchment. Once this erosion process has been initiated in these locations, a greater volume of sediment is available for erosion and, because of the close spatial proximity of these erosion sites to waterways, that eroded sediment is likely to be transported efficiently to waterways. Alluvial gully erosion has been reported in other catchments in northern Australia (Brooks et al., 2013; Daley et al., 2021; Shellberg et al., 2016) and the results presented here are likely to be relevant to sediment budgets created for those catchments, and any extensions of such sediment budgets which estimate the sediment retention ecosystem service for use in SEEA EA.

While the above are important considerations for how the sediment retention ecosystem service is represented, this ecosystem service may not have a high monetary value in the Mitchell catchment. The loss of productive agricultural and grazing land due to erosion will impose a cost on local landowners; however, although gullying is quite extensive in some locations it still affects only a minor fraction of the land area of the large grazing properties in the catchment.

Furthermore, due to the small population affected by reduced water quality downstream sediment erosion will not have major direct adverse implications, for example on drinking water supply (UN, 2021b). Nevertheless, despite the small population in the catchment, there are several significant human uses that may be adversely affected by loss of the sediment retention ecosystem service. For example the catchment supports a valuable commercial barramundi fishery and increases in sediment load may affect important habitats and subsequent recruitment into the fishery (Bayliss et al., 2014; Pollino et al., 2018). In addition, the local Indigenous population gains significant benefits from being able to source food using traditional methods in the catchment, and any changes to water quality which affect the viability of Indigenous food sources would have major implications for their wellbeing (Jackson et al., 2011).

The results presented here have allowed an exploration of the implications of using a simple approach to model the sediment retention ecosystem service, and to identify locations within the Mitchell catchment that supply this ecosystem service. The Mitchell catchment has high ecological value (Pollino et al., 2018), and has experienced a relatively high level of erosion in response to a relatively modest historical human impact when compared to catchments in southern Australia. In this sense it provides an ideal case study to examine the ability of SEEA EA ecosystem service modelling tools to estimate the soil retention service, and guide investment in ecosystem restoration. The results indicate that a simplified representation of erosion processes would lead to misinformed management responses (e.g. interventions to reduce hillslope erosion) in inappropriate locations (e.g. the upper catchment). These limitations are highly relevant if InVEST is to be used to inform SEEA EA applications to manage both ends of the spectrum of human modifications of ecosystems - the few remaining areas with minimal human influence and high ecological value and/or areas of high human modification which are supplying substantial ecosystem service flows to large population centres downstream.

### Wider implications of sediment retention modelling

The large spatial areas and long temporal scales relevant to the transport of sediment from erosion sources to a catchment outlet, and the substantial variability through which these processes occur, present major challenges to both measurement and modelling (Walling & Collins, 2008). The sediment budget is a means to assemble process understanding and explore geomorphic system behaviour, explore scenarios, generate hypotheses, and check for inconsistencies against multiple lines of evidence in order to target further research and data collection (Rustomji et al., 2008; Rustomji et al., 2010; Silberstein, 2006; Walling & Collins, 2008). While the sediment budget approach holds promise for improving our understanding of both on- and off-site implications of soil erosion, substantial challenges remain in assembling the data and information required for creating reliable sediment budgets (Walling & Collins, 2008).

The sediment budget methodology and the importance of using a calibrated and validated model for ecosystem service valuation is acknowledged in the InVEST documentation (Sharp et al., 2018), and research is progressing on refining the hillslope erosion and sediment delivery approach that InVEST uses currently (Hamel et al., 2015; Hamel et al., 2017). The results presented here allowed the difference in ecosystem service flow (in physical terms) and service providing areas to be compared between a model version which represented hillslope erosion and sediment delivery in isolation, a model version which included other erosion and deposition processes, and between an initial model and a final model calibrated against available empirical data. The results demonstrated that if hillslope erosion was indeed the dominant erosion process at a location, and if relative magnitude, rather than the absolute magnitude, of erosion is the focus, then RUSLE may not lead to excessive errors. However if hillslope erosion is not the dominant process, as was the case in the Mitchell, a solely RUSLE-based approach may identify incorrect locations as supplying the majority of the sediment retention ecosystem service, and may therefore lead to incorrect interventions being suggested to increase the flow of this service. For example, a RUSLE-only approach would identify hillslopes in the upper Mitchell catchment as being the main focus for management intervention to increase supply of the sediment retention service. However when using the calibrated sediment budget model, alluvial sections of the mid to lower catchment were identified as the main focus for intervention to increase sediment retention (see the results from the pre-European condition sediment budget).

One of the goals of SEEA EA is to compile information on ecosystems’ contributions to the economy and society to guide decisions about investment in ecosystem protection and rehabilitation (UN, 2021b). This paper has demonstrated the need to review model predictions alongside relevant local data and in close consultation with relevant physical process experts (for example hydrologists and/or geomorphologists). If estimates of the supply of the sediment retention ecosystem service across a catchment were informed by InVEST alone and hillslope erosion was not the dominant erosion process contributing to suspended sediment load in a region, it is possible that the limited funds available may be misallocated, both from a management response perspective and from a spatial prioritisation perspective. This has the potential to reduce key stakeholders’ and decision makers’ willingness to apply findings from SEEA EA (Guerry et al., 2015; Naeem et al., 2015). This is an ongoing challenge as the sediment retention ecosystem service can supply considerable value to society. However, the research and data required to robustly identify the most important erosion processes and their links to suspended sediment yield at the catchment outlet are unlikely to be present in many locations.

The results summarised here for the Mitchell catchment are valuable for assessing the potential implications of using a simplified representation of this ecosystem service.

## Conclusion

Management of soil erosion and water quality is highly relevant to land and water sustainability and many of the United Nations Sustainable Development Goals. However, challenges remain for measuring and modelling these processes which occur over large areas and long-time scales, and experience substantial variability in rates. It is clear that this management problem needs to be addressed, however the results presented for the Mitchell catchment demonstrate that practitioners must exercise caution when using estimates of the sediment retention ecosystem service flow which have not been calibrated and validated against locally collected empirical data. Further research is needed on approaches for improving estimates of this ecosystem service in data poor regions.

## References

Amundson, R., Berhe, A. A., Hopmans, J. W., Olson, C., Sztein, A. E., & Sparks, D. L. (2015). Soil and human security in the 21st century. Science, 348(6235), 1261071.

Armston, J. D., Danaher, T. J., & Collett, L. J. (2004). A regression approach for mapping woody foliage projective cover in Queensland with Landsat data. Paper presented at the Proceedings of the 12th Australasian Remote Sensing and Photogrammetry Conference, Fremantle, Australia.

Bayliss, P., Buckworth, R., & Dichmont, C. (2014). Assessing the water needs of fisheries and ecological values in the Gulf of Carpentaria. Final Report prepared for the Queensland Department of Natural Resources and Mines (DNRM), CSIRO, Australia

Borselli, L., Cassi, P., & Torri, D. (2008). Prolegomena to sediment and flow connectivity in the landscape: A GIS and field numerical assessment. Catena, 75(3), 268–277.

Brierley, G. J., & Fryirs, K. A. (2005). Geomorphology and river management: Applications of the river styles framework. Malden, MA: Blackwell Publishing.

Brooks, A., Spencer, J., Borombovits, D., Pietsch, T., & Olley, J. (2014). Measured Hillslope Erosion Rates in the Wet-Dry Tropics of Cape York, Northern Australia Part 2: Rusle-Based Modeling Significantly Over-Predicts Hillslope Sediment Production. Catena, 122, 1–17.

Brooks, A. P., Shellberg, J. G., Knight, J., & Spencer, J. (2009). Alluvial gully erosion: An example from the Mitchell fluvial megafan, Queensland, Australia. Earth Surface Processes and Landforms, 34(14), 1951–1969.

Brooks, A. P., Spencer, J., Olley, J., Pietsch, T., Borombovits, D., Curwen, G., … Bourgeault, A. (2013). An empirically-based sediment budget for the Normanby Basin: Sediment sources, sinks, and drivers on the Cape York Savannah. Griffith University, Brisbane

Brooks, A. P., Spencer, J., Shellberg, J. G., Knight, J., & Lymburner, L. (2008). Using remote sensing to quantify sediment budget components in a large tropical river - Mitchell River, Gulf of Carpentaria. IAHS PUBLICATION(325), 225–236.

Caitcheon, G., Hancock, G., Leslie, C., & Olley, J. (2011). Regional scale sediment and nutrient budgets - Sediment tracing report on TRaCK Project 4.2. Technical report, Tropical Rivers and Coastal Knowledge,

Caitcheon, G. G., Olley, J. M., Pantus, F., Hancock, G., & Leslie, C. (2012). The dominant erosion processes supplying fine sediment to three major rivers in tropical Australia, the Daly (NT), Mitchell (Qld) and Flinders (Qld) River. Geomorphology, 151–152, 188-195.

Collins, A. L., & Walling, D. E. (2004). Documenting catchment suspended sediment sources: Problems, approaches and prospects. Progress in Physical Geography, 28(2), 159–196.

Daley, J., Stout, J. C., Curwen, G., Brooks., A. P., Spencer., J., Pietsch., T., & Thwaites, R. (2021). Development and application of automated tools for high resolution gully mapping and classification from lidar data. Report to the National Environmental Science Program, Reef and Rainforest Research Centre Limited, Cairns

Fryirs, K. (2013). (Dis)Connectivity in catchment sediment cascades: A fresh look at the sediment delivery problem. Earth Surface Processes and Landforms, 38, 30–46.

Gallant, J. C., & Dowling, T. I. (2003). A multiresolution index of valley bottom flatness for mapping depositional areas. Water Resources Research, 39(12), WR001426.

Guerry, A. D., Polasky, S., Lubchencof, J., Chaplin-Kramerb, R., Daily, G. C., Griffin, R., … Viraw, B. (2015). Natural capital and ecosystem services informing decisions: From promise to practice. Proceedings of the National Academy of Sciences of the United States of America, 112(24), 7348–7355.

Hamel, P., Chaplin-Kramer, R., Sim, S., & Mueller, C. (2015). A new approach to modeling the sediment retention service (InVEST 3.0): Case study of the Cape Fear catchment, North Carolina, USA. Science of the Total Environment, 524-525, 166–177.

Hamel, P., Sharp, R., Dennedy-Frank, P. J., Falinski, K., Auerbach, D. A., Sanchez-Canales, M., & Dennedy-Frank, P. J. (2017). Sediment delivery modeling in practice: Comparing the effects of watershed characteristics and data resolution across hydroclimatic regions. Science of the Total Environment, 580, 1381–1388.

Hughes, A., Prosser, P., Stevenson, J., Scott, A., Lu, H., Gallant, J., & Moran, C. (2001). Gully erosion mapping for the National Land and Water Audit (Technical Report 26/01). CSIRO Land and Water, Canberra

Jackson, S., Finn, M., Woodward, E., & Featherston, P. (2011). Indigenous socio-economic values and river flows. CSIRO Ecosystem Sciences, Darwin, NT

Keeler, B. L., Polasky, S., Brauman, K. A., Johnson, K. A., Finlay, J. C., O’Neill, A., … Dalzell, B. (2012). Linking water quality and well-being for improved assessment and valuation of ecosystem services. Proceedings of the National Academy of Sciences, 109(45), 18619–18624.

Keesstra, S. D., Bouma, J., Wallinga, J., Tittonell, P., Smith, P., Cerdà, A., … Fresco, L. O. (2016). The significance of soils and soil science towards realization of the United Nations Sustainable Development Goals. Soil, 2(2), 111–128.

Lu, H., Prosser, I. P., Moran, C. J., Gallant, J. C., Priestley, G., & Stevenson, J. G. (2003). Predicting sheetwash and rill erosion over the Australian continent. Australian Journal of Soil Research, 41, 1037–1062.

McCloskey, G. L., Baheerathan, R., Dougall, C., Ellis, R., Bennett, F. R., Waters, D., … Askildsen, M. (2021). Modelled estimates of fine sediment and particulate nutrients delivered from the Great Barrier Reef catchments. Marine pollution bulletin, 165, 112163.

Naeem, S., Ingram, J. C., Varga, A., Agardy, T., Barten, P., Bennett, G., … Wunder, S. (2015). Get the science right when paying for nature’s services. Science, 347(6227), 1206–1207.

Olley, J., Brooks, A., Spencer, J., Pietsch, T., & Borombovits, D. (2013). Subsoil erosion dominates the supply of fine sediment to rivers draining into Princess Charlotte Bay, Australia. Journal of Environmental Radioactivity, 124(22), 121–129.

Olley, J. M., Murray, A. S., Mackenzie, D. H., & Edwards, K. (1993). Identifying sediment sources in a gullied catchment using natural and anthropogenic radioactivity. Water Resources Research, 29(4), 1037.

Petheram, C., Watson, I., Bruce, C., & Chilcott, C. (2018). Water resource assessment for the Mitchell catchment. A report to the Australian Government from the CSIRO Northern Australia Water Resource Assessment, part of the National Water Infrastructure Development Fund: Water Resource Assessments, CSIRO, Australia

Pickup, G., & Marks, A. (2001). Identification of floodplains and estimation of floodplain flow velocities for sediment transport modelling. Technical Report 14/01, CSIRO Land and Water, Canberra

PMC. (2012). Australia in the Asian century. White Paper, Department of Prime Minister and Cabinet, Australia

Pollino, C. A., Barber, E., Buckworth, R., Cadiegues, M., Cook, G., Deng, A., … Turschwell, M. (2018). Synthesis of knowledge to support the assessment of impacts of water resource development to ecological assets in northern Australia: Asset analysis. A technical report to the Australian Government from the CSIRO Northern Australia Water Resource Assessment, part of the National Water Infrastructure Development Fund: Water Resource Assessments, CSIRO, Australia

Prosser, I., Rustomji, P., Young, P., Moran, C., & Hughes, A. (2001a). Constructing river basin sediment budgets for the National Land and Water Audit. CSIRO Land and Water, Report No. 15/01, Canberra

Prosser, I. P., Rutherfurd, I. D., Olley, J. M., Young, W. J., Wallbrink, P. J., & Moran, C. J. (2001b). Large-scale patterns of erosion and sediment transport in river networks, with examples from Australia. Marine and Freshwater Research, 52, 81–99.

Qld Government. (2020). Types of erosion. Queensland Government. Retrieved from https://www.qld.gov.au/environment/land/management/soil/erosion/types

Renard, K. G., Foster, G. A., Weesies, D. K., McCool, D. K., & Yoder, D. C. (1997). Predicting soil erosion by water: A guide to conservation planning with the Revised Universal Soil Loss Equation. Agriculture Handbook 703, United States Department of Agriculture, Washington DC

Rustomji, P. (2010). A statistical analysis of flood hydrology and bankfull discharge for the Mitchell River catchment, Queensland, Australia. Water for a Healthy Country Report, CSIRO, Canberra

Rustomji, P., Caitcheon, G., & Hairsine, P. B. (2008). Combining a spatial model with geochemical tracers and river station data to construct a catchment sediment budget. Water Resources Research, 44(W01422).

Rustomji, P., Shellberg, J., Brooks, A., Spencer, J., & Caitcheon, G. (2010). A catchment sediment and nutrient budget for the Mitchell River, Queensland. A report to the Tropical Rivers and Coastal Knowledge (TRaCK) Research Program, CSIRO Water for a Healthy Country National Research Flagship,

Schumm, S. A., & Lichty, R. W. (1965). Time, space and causality in geomorphology. American Journal of Science, 263, 110–119.

Sharp, R., Tallis, H. T., Ricketts, T., Guerry, A. D., Wood, S. A., Chaplin-Kramer, R., … Douglass, J. (2018). InVEST 3.5.0 User’s Guide. The Natural Capital Project, Stanford University, University of Minnesota, The Nature Conservancy, and World Wildlife Fund,

Shellberg, J., Brooks, A., & Spencer, J. (2010a, 1-6 August 2010.). Land-use change from indigenous management to cattle grazing initiates the gullying of alluvial soils in northern Australia. Paper presented at the 19th World Congress of Soil Science: Soil solutions for a changing world, Brisbane, Australia.

Shellberg, J. G., Brooks, A. P., & Rose, C. W. (2013a). Sediment production and yield from an alluvial gully in Northern Queensland, Australia. Earth Surface Processes and Landforms, 38(15), 1765–1778. Retrieved from http://onlinelibrary.wiley.com/store/10.1002/esp.3414/asset/esp3414.pdf?v=1&t=imr8xt8i&s=db6722f8445a266841b038cc50aa86f244430d88

Shellberg, J. G., Brooks, A. P., Spencer, J., Knight, J., & Pietsch, T. (2010b). Alluvial gully erosion rates and processes in Northern Queensland: an example from the Mitchell River fluvial megafan.

Produced for The Caring for Our Country (CfoC) Initiative, Australian Rivers Institute, Griffith University,

Shellberg, J. G., Brooks, A. P., Spencer, J., & Ward, D. (2013b). The hydrogeomorphic influences on alluvial gully erosion along the Mitchell River fluvial megafan. Hydrological Processes, 27(7), 1086–1104.

Shellberg, J. G., Spencer, J., Brooks, A. P., & Pietsch, T. J. (2016). Degradation of the Mitchell River fluvial megafan by alluvial gully erosion increased by post-European land use change, Queensland, Australia. Geomorphology, 266, 105–120.

Silberstein, R. P. (2006). Hydrological models are so good, do we still need data? Environmental Modelling and Software, 21, 1340–1352.

Trimble, S. W. (1983). A sediment budget for Coon Creek basin in the Driftless Area, Wisconsin, 1853-1977. American Journal of Science, 283, 454–474.

UN. (2021a). Guidelines on biophysical modelling for ecosystem accounting – Version 2.0.

United Nations, UN. (2021b). System of Environmental-Economic Accounting - Ecosystem Accounting. Statistics Division, United Nations,

Wallbrink, P. J., & Murray, A. S. (1993). Use of fallout radionuclides as indicators of erosion processes. Hydrological Processes, 7(3), 297–304.

Walling, D. E., & Collins, A. L. (2005). Suspended sediment sources in British rivers. IAHS PUBLICATION(291), 123–133.

Walling, D. E., & Collins, A. L. (2008). The catchment sediment budget as a management tool. Environmental Science and Policy, 11(2), 136–143.

Wasson, R. J. (1994). Annual and decadal variation of sediment yield in Australia, and some global comparisons. Paper presented at the Variability in stream erosion and sediment transport (Proceedings of the Canberra Symposium), Canberra.

Wilkinson, S., Henderson, A., Chen, Y., & Sherman, B. (2008). SedNet User Guide. Client Report, CSIRO Land and Water, Canberra

Wilkinson, S. N., Dougall, C., Kinsey-Henderson, A. E., Searle, R. D., Ellis, R. J., & Bartley, R. (2014). Development of a time-stepping sediment budget model for assessing land use impacts in large river basins. Science of the Total Environment, 468-469, 1210–1224.

Wilkinson, S. N., Prosser, I. P., Rustomji, P., & Read, A. M. (2009). Modelling and testing spatially distributed sediment budgets to relate erosion processes to sediment yields. Environmental Modelling and Software, 24, 489–501.

Wilkinson, S. N., Young, W. J., & De Rose, R. C. (2006). Regionalising mean annual flow and daily flow variability for bain-scale sediment and nutrient modelling. Hydrological Processes, 20, 2769–2786.

Wishcheimer, W., & Smith, D. D. (1978). Predicting rainfall erosion lossess: A guide to conservation planning (Agriculture Handbook No. 537). US Department of Agriculture, Washington

